# Robust, Universal Tree Balance Indices

**DOI:** 10.1101/2021.08.25.457695

**Authors:** Jeanne Lemant, Cécile Le Sueur, Veselin Manojlović, Robert Noble

## Abstract

Balance indices that quantify the symmetry of branching events and the compactness of trees are widely used to compare evolutionary processes or tree-generating algorithms. Yet existing indices have important shortcomings, including that they are unsuited to the tree types commonly used to describe the evolution of tumours, microbial populations, and cell lines. The contributions of this article are twofold. First, we define a new class of robust, universal tree balance indices. These indices take a form similar to Colless’ index but account for node sizes, are defined for trees with any degree distribution, and enable more meaningful comparison of trees with different numbers of leaves. Second, we show that for bifurcating and all other full m-ary cladograms (in which every internal node has the same out-degree), one such Colless-like index is equivalent to the normalised reciprocal of Sackin’s index. Hence we both unify and generalise the two most popular existing tree balance indices. Our indices are intrinsically normalised and can be computed in linear time. We conclude that these more widely applicable indices have potential to supersede those in current use.

---

Tree balance indices – most notably those credited to Sackin (1972) and Colless (1982) – are widely used to describe speciation processes, compare cladograms, and assert the correctness of tree reconstruction methods (Shao and Sokal, 1990; Mooers and Heard, 1997). These indices have recently been introduced to oncology (Chkhaidze et al., 2019; Scott et al., 2020) because methods for determining and classifying modes of tumour evolution have clinical value (Maley et al., 2017). A problem here is that the trees that best describe tumour evolution are clone trees in which node sizes are informative and which frequently contain linear sections; indeed, developing methods to distinguish linear from branching tumour evolution is an important area of ongoing research (Davis et al., 2017). Existing tree balance indices are unsuited to these topologies and take no account of node size. Moreover, even when applied only to bifurcating cladograms, existing indices are unreliable for comparing trees with different numbers of leaves.

Here we develop a new class of robust, universal tree balance indices. Our definitions not only extend the tree balance concept and open up new applications but also unify the two main approaches to quantifying balance as proposed by Sackin and Colless. We describe several general advantages of our indices compared to those in current use.

## Materials and Methods

### Rooted trees

We consider exclusively rooted trees in which all edges are oriented away from the root (which will be topmost in our figures). This orientation defines a natural order on the tree, from top to bottom: all edges are assumed to extend from the root to the other *internal nodes* and finally to the terminal nodes or *leaves*. The *out-degree* of a node *i*, written *d*^+^(*i*), is the number of direct descendants, ignoring any descendant branches in which all nodes have zero size. Internal nodes have out-degree at least one, whereas leaves have out-degree zero.

Some tree types have particular names. A *caterpillar tree* (Fig. 1a) is a bifurcating tree in which each internal node has one leaf. A *fully symmetric* tree (Fig. 1b) is such that every internal node with the same depth has the same degree or, equivalently, for each internal node *i* all the subtrees rooted at *i* are identical. A *star tree* (Fig. 1c) is a tree whose leaves are all attached to the root, which is the only internal node.

**Figure 1:**
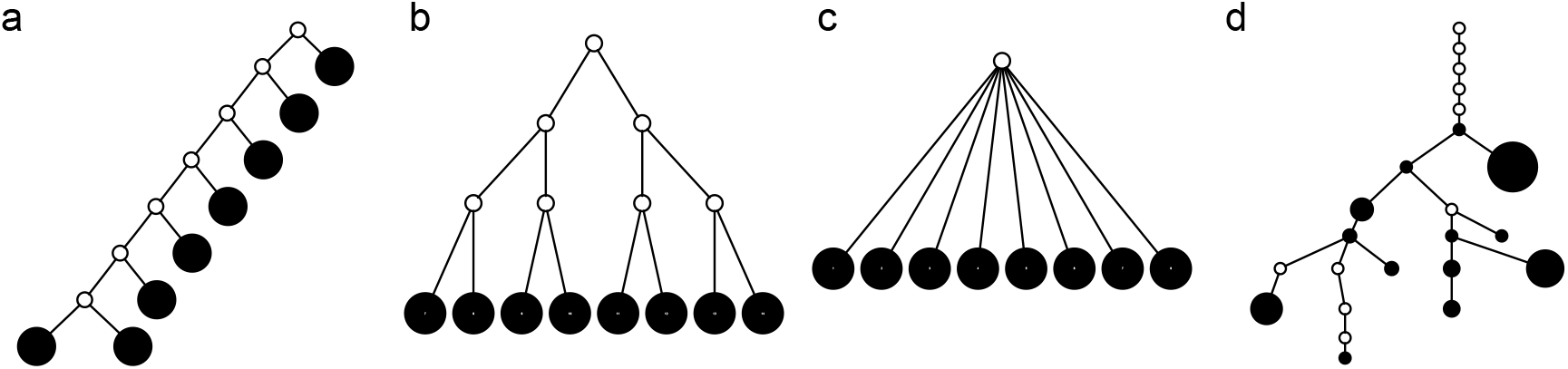
Contrasting trees. **a**: Caterpillar tree with *I*_*S*_ = 35, *I*_*S,norm*_ = 1, *I*_*C*_ = 21, *I*_*C,norm*_ = 1, *I*_Φ_ = 56, *I*_Φ,*norm*_ = 1. **b**: Fully symmetric bifurcating tree with *I*_*S*_ = 24, *I*_*S,norm*_ ≈ 0.59, *I*_*C*_ = *I*_*C,norm*_ = 0, *I*_Φ_ = 16, *I*_Φ,*norm*_ ≈ 0.29. **c**: Star tree with *I*_*S*_ = 8, *I*_*S,norm*_ = 0, *I*_*C*_ and *I*_*C,norm*_ undefined, *I*_Φ_ = *I*_Φ,*norm*_ = 0. **d**: Clone tree of the lung tumour CRUK0065 in the TRACERx cohort (Jamal-Hanjani et al., 2017). In the clone tree, nodes represented by empty circles correspond to extinct clones, and the diameters of other nodes are proportional to the corresponding clone population sizes.

### Cladograms, species trees and clone trees

Cladograms are trees that represent relationships between extant biological taxa (leaves) via edges linking them to hypothetical extinct ancestors (internal nodes). A common conception is that only bifurcating cladograms can be considered fully resolved and linear parts are inadmissible. However, linear sections in cladograms are appropriate for representing anagenesis (in which a descendant replaces its ancestor), while budding (in which an ancestor produces a descendant and remains extant) can give rise to cladogram nodes with out-degree greater than two (Podani, 2013). An extant ancestor is represented in a cladogram by a leaf stemming from the internal ancestor node (so the two nodes represent the same taxon).

An alternative way to represent extant ancestors is as internal nodes (like in a genealogy with overlapping generations). Such diagrams are known to organismal biologists as species trees and to oncologists as clone trees. In a clone tree, each node represents a clone (a set of cells that share alterations of interest due to common descent) and edges represent the chronology of alterations. Clone tree nodes can have any out-degree, including *d*^+^ = 1, and each node – including internal nodes – can be associated with a non-negative size, related to the clone population size at the time of observation (as in Figure 1d). The size of a tree or subtree can then be defined as the sum of its node sizes.

When nodes are associated with sizes, the addition or removal of even vanishingly small terminal branches can change leaves into internal nodes or vice versa and so substantially change the value of existing tree balance indices. This behaviour is unsatisfactory because these small branches typically represent either newly-created types that have yet to experience evolutionary forces or types on the verge of extinction, and in either case their relative sizes convey negligible information about the mode of evolution. Data sets may also omit rare types due to sampling error or because genetic sequencing methods have imperfect sensitivity (Turajlic et al., 2018).

The change due to the addition of terminal nodes is greater when the tree is a cladogram rather than a species or clone tree. For example, when a three-node, two-leaf tree (Fig. 2a) is augmented by adding a node *j* to a leaf *i* (Fig. 2b), the three original nodes retain their positions in the species or clone tree (middle column of Figure 2), but in the cladogram (right column) node *i* becomes two nodes (*i*_1_ and *i*_2_), the larger of which is now further from the root. As the size of the new node *j* is continuously reduced to zero, the species or clone tree changes continuously, whereas the cladogram undergoes an abrupt change of topology when the size of node *j* reaches zero. We conclude that the species or clone tree representation is more robust than the cladogram representation in the general case in which nodes are associated with sizes and ancestors can be extant. Also an index that accounts for non-zero internal node sizes can be made more robust than one that does not.

**Figure 2:**
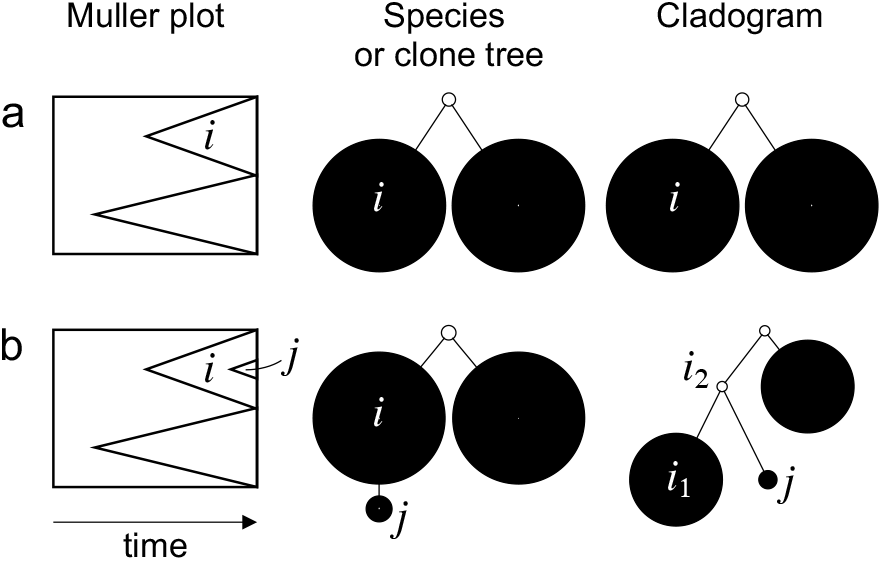
Muller plots (left column), species or clone trees (middle column), and cladograms (right column) representing evolution by splitting only (a) and both splitting and budding (b). Nodes represented by empty circles correspond to extinct types.

### Existing tree balance indices

The most widely used tree balance indices are in fact imbalance indices, such that more balanced trees are assigned smaller values. These indices were introduced to study cladograms and take no account of node size. The most popular are Sackin’s index and Colless’ index.

#### Sackin’s index

Let *T* be a tree with set of leaves *L*(*T*). For a leaf *l* ∈ *L*(*T*), let *ν*_*l*_ be the number of internal nodes between *l* and the root, which is included in the count. Then the index credited to Sackin (1972) is

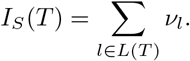

For two bifurcating trees on the same number of leaves, a less balanced tree has higher values of *ν* as the tree is in a sense less compact (compare trees **a** and **b** in Figure 1).

Since the value tends to increase with the number of nodes, Shao and Sokal (1990) proposed normalising *I*_*S*_ with respect to trees on *n* > 2 leaves by subtracting its minimum possible value for such trees and then dividing by the difference between the maximum and minimum possible values. The minimal *I*_*S*_ is reached on the star tree, such as tree **c** in Figure 1, and hence min_*n*_(*I*_*S*_) = *n*. The maximum is attained on the caterpillar tree, such as tree **a**:

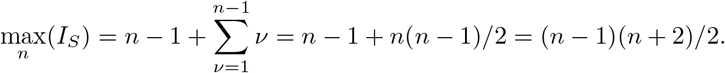

The normalised index is then

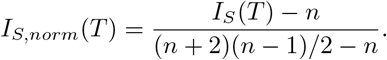

This normalised index is not very satisfactory as a balance index because it fails to capture an intuitive notion of balance. For example, it is not obvious why fully symmetric tree **b** should be considered less balanced than star tree **c** in Figure 1, yet its *I*_*S,norm*_ value is much larger. To address this issue, Shao and Sokal (1990) further suggested normalising *I*_*S*_ relative to its extremal values among trees with the same number of internal nodes as well as the same number of leaves. But even then the index remains unreliable for comparing trees with different numbers of leaves. For example, the index is 1 for every caterpillar tree, yet long caterpillar trees are intuitively less balanced than short ones. The conventional *I*_*S*_ normalisations are not defined for trees containing linear parts. Moreover, since *I*_*S*_ doesn’t account for node size, it is highly sensitive to the addition or removal of relatively tiny terminal branches. Hence Sackin’s index is neither universal nor robust.

#### Colless’ index

For an internal node *i* of a bifurcating tree *T*, define 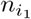 as the number of leaves of the left branch of the subtree rooted at *i*, and 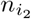 as the number of leaves of the right branch. Then the index defined by Colless (1982) is

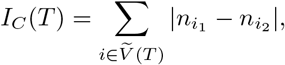

where 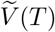 is the set of internal nodes. The index can be normalised for the set of trees on *n* > 2 leaves by dividing by its maximal value, 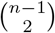 which is reached on the caterpillar tree (as in Figure 1a).

To generalise Colless’ index to multifurcating trees, Mir et al. (2018) recently introduced a family of Colless-like balance indices, including *I*_*C*_ as a special case. Each of these indices 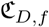 is determined by a weight function *f*, which assigns a size to each subtree as a function of its out-degree, and a dissimilarity function *D*. By definition of *D*, Colless-like indices are zero if and only if each internal node divides its descendants into subtrees of equal size according to *f*. But since these indices are normalised by dividing by the maximal value for trees on the same number of leaves, they are unreliable for comparing trees with different numbers of leaves. In common with Sackin’s index, the total cophenetic index *I*_Φ_ (Mir et al., 2013) (see Appendix), and other existing indices, the Colless-like indices so far defined are neither universal nor robust.

### Desirable properties of a universal, robust tree balance index

Our aim is to derive a tree balance index *J* that is useful for classifying and comparing rooted trees that can have any distributions of node degrees and node sizes. Here we specify five desirable properties that such an index should have. The first two axioms relate to extrema and universality, in the sense of an index being defined for trees with any degree distribution. The other three axioms are concerned with robustness and are relevant only when nodes can have arbitrary sizes.

Conventionally, a tree is considered maximally balanced only if every internal node splits its descendants into subtrees on the same number of leaves (Shao and Sokal, 1990). We generalise this concept by requiring that every internal node splits its descendants into at least two subtrees of equal size, as in Figure 3a. We term this the *equal splits* property. We then set necessary and sufficient conditions for maximal balance:

#### Axiom 0.1 (Maximum value).

*J*(*T*) ≤ 1 *for all trees T, and J* (*T*) = 1 *only if T has equal splits. Furthermore, if T has equal splits and every internal node of T has null size (or equivalently represents an extinct taxon) then J* (*T*) = 1.

Another convention is that narrow trees with relatively many internal nodes are considered highly imbalanced. Linear trees (that is, trees in which every node *i* has *d*^+^(*i*) ≤ 1, as in Figure 3b) are even narrower than caterpillar trees. Also the most unequal binary split is one that assigns all descendants to one branch and none to the other. Hence our second desirable property:

#### Axiom 0.2 (Minimum value).

*J* ≥ (*T*) 0 *for all trees T, and J* (*T*) = 0 *if and only if T is a linear tree.*

Our third desirable property is that our index should be insensitive to the presence of uninformative terminal branches:

#### Axiom 0.3 (Leaf limit).

*Let T be a tree with finitely many nodes and l be a leaf of T. Suppose we create a new tree T* ′ *by adding to T a subtree T_l_ with finitely many nodes, rooted at l. As the size of T_l_ excluding its root approaches zero, so J* (*T′*) → *J*(*T*).

Our fourth desirable property ensures that a linear section of a tree is regarded as a maximally unequal split:

#### Axiom 0.4 (Linear limit).

*Let j be a node of a tree T with d*^+^(*j*) = 1*. Suppose we create a new tree T* ′ *by adding to T a subtree with finitely many nodes, rooted at j. As the size of T_j_ excluding its root approaches zero, so J* (*T′*) → *J*(*T*).

Lastly, we require continuity with respect to varying node size:

#### Axiom 0.5 (Continuity).

*If the population of any node of any tree T varies continuously in* ℝ_>0_, *then J*(*T*) *varies continuously.*

The wording of Axiom 0.1 raises an important question: Can trees with non-zero-sized internal nodes be considered maximally balanced? The following proposition provides the answer.

#### Proposition 0.6.

*Axioms 0.3 and 0.4 each imply that equal splits are not sufficient for maximal balance.*

*Proof.* Suppose that equal splits are sufficient for maximal balance. First consider a one-node tree *T*. If we add a vanishingly small linear subtree to *T* then the new tree *T′* will have *J*(*T′*) = 0. But if we instead add two vanishingly small subtrees of equal size to *T* then we obtain *J*(*T′*) = 1. This implies that whatever value we assign to *J* (*T*), we cannot satisfy Axiom 0.3. Second, consider a linear tree *T* in which the sum of the non-root node sizes is *δ*. Then *J*(*T*) = 0. But if we add another subtree to the root, also of size *δ*, then the new tree *T′* will have *J*(*T′*) = 1, even as *δ* → 0. This contradicts Axiom 0.4.

**Figure 3:**
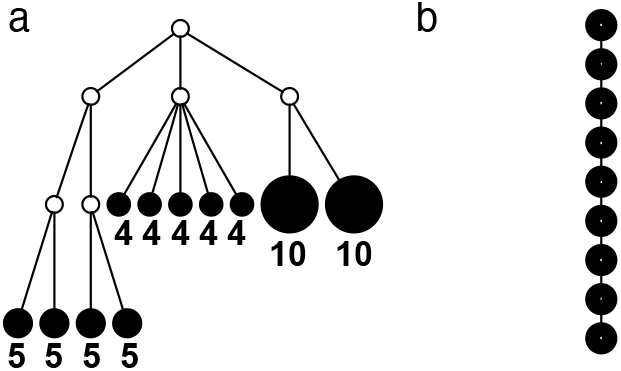
Trees with extremal *J* values. Numbers shown below nodes are node sizes. Empty nodes have null size. **a**: A tree in which each internal node has null size and splits its descendants into subtrees of equal size, and hence *J* = 1. This tree can be considered balanced only according to an index that accounts for node size. **b**: A linear tree, for which *J* = 0.

We therefore face a choice: either weaken Axioms 0.3 and 0.4 or accept that equal splits are not sufficient for maximal balance. We choose the second option (and as a corollary obtain *J* = 0 for the single-node tree) because we want our indices to be not only universal but also highly robust when applied to real, imperfect data. We will further argue that this choice is appropriate from a biological viewpoint and is consistent with the ideas underlying previous tree balance indices.

## Results

### General definition of universal, robust tree balance indices

Before defining a new class of balance indices we need to introduce some more notation. For a tree *T*, we will use *V*(*T*) to denote the set of all nodes of *T*, which we will abbreviate to *V* when the identity of the tree is unambiguous. Let *f*(*v*) ≥ 0 denote the size of node *v* (not necessarily a function of the out-degree). Then *S*_*i*_ denotes the size of the subtree *T*_*i*_ rooted at *i*, and 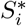 is the size of *T*_*i*_ excluding its root:

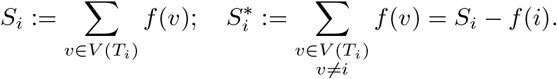

We will use 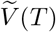 or simply 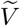 to denote the set of all internal nodes such that 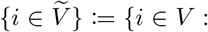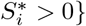

We then introduce three continuous functions of subtree sizes:

- An *importance* factor *g* : ℝ_>0_ → ℝ_>0_ with *g*(*x*) → 0 as *x* → 0;
- A *non-root dominance* factor *h* : ℝ_>0_ × ℝ_>0_ → (0, 1] with *h*(*x*_1_, *x*_2_) → 0 as *x*_1_ → 0, and *h*(*x*_1_, *x*_2_) = 1 if and only if *x*_1_ = *x*_2_;
- A *balance score W* that assigns *W*_*i*_ ∈ [0, 1] to each internal node *i* such that *W*_*i*_ = 0 if and only if *d*^+^(*i*) = 1, and *W*_*i*_ = 1 if and only if *i* splits its descendants into at least two equally sized subtrees.

To allow us to define *W* more rigourously, let 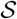 denote the set of vectors with positive components that sum to unity:

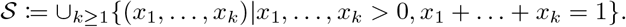

Then 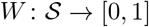 is such that, for all 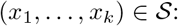

- For every permutation *π*, *W*(*x*_1_, …, *x*_*k*_) = *W* (*x*_*π*(1)_, …, *x*_*π*(*k*)_);
- *W*(*x*_1_, …, *x*_*k*_) = 1 if and only if *k* > 1 and *x*_1_ = … = *x*_*k*_;
- *W* = 0 if and only if max(*x*_1_, …, *x*_*k*_) = 1;
- *W* is a continuous function with respect to each of its arguments.

We then define a balance index in terms of subtree sizes as

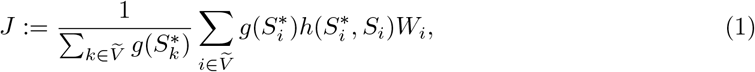

where 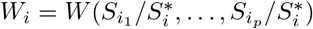 and *i*_1_, …, *i*_*p*_ are the children of node *i*. A short proof that this type of index satisfies our five axioms for robustness and universality is presented in the Appendix.

### Interpretation of factors *W*, *g* and *h*

The balance score *W* in our general definition (Equation 1) measures the extent to which an internal node splits its descendants into equally sized subtrees. The importance factor *g* assigns more weight to nodes that are the roots of large subtrees. In biological terms, this means giving more weight to types that have more descendants. The continuous function *h* quantifies the extent to which a node should be considered a leaf (which doesn’t contribute to determining tree balance in Colless-like indices) as opposed to an internal node (which does). From a biological point of view, nodes that are large relative to their descendants represent extant populations whose evolutionary fate remains largely undetermined.

Factors *g* and *h* together ensure that our indices consider a tree imbalanced unless there is strong evidence to the contrary. For example, the one-node tree provides no evidence and is considered maximally imbalanced. The inclusion of *h* also means that a tree can achieve maximal balance only if all its internal nodes have zero size, which is equivalent to all ancestors being extinct, as in a cladogram. This requirement can be removed simply by omitting *h* from the definition, but then, as per Proposition 0.6, our robustness Axioms 0.3 and 0.4 will not be satisfied.

Sackin’s and Colless’ indices similarly assign more weight to nodes that have more descendant leaves or are closer to the root. As Mooers and Heard (1997) have remarked, it is reasonable to put more weight on nodes deeper within the tree because “those nodes are the most informative, as the subclades they define are older and therefore sample longer periods of evolutionary time.”

### A specific index based on the Shannon entropy

In defining a specific index, we start by opting for the simplest choices of importance and non-root dominance factors:

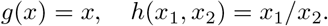

The role of the balance score function *W* is to quantify the extent to which a set of objects (specifically subtrees) have equal size. A well-known index that satisfies the necessary conditions is the normalised Shannon entropy.

Assume a population is partitioned into *n* ∈ ℕ types, with each type *i* accounting for a proportion *p*_*i*_. Then the Shannon entropy with base *b* is defined as 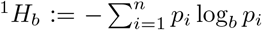. If all types have equal frequencies *p*_*i*_ = 1/*n* then ^1^*H*_*b*_ = log_*b*_*n*. If the types have unequal sizes then ^1^*H*_*b*_ < log_*b*_*n*. And if the abundance is mostly concentrated on one type *j*, such that *p*_*j*_ → 1, then ^1^*H*_*b*_ → 0.

Let *C*(*i*) denote the set of children (immediate descendants) of a node *i*, and for *j* ∈ *C*(*i*) let 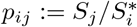 denote the relative size of subtree *T*_*j*_ compared to all subtrees attached to *i*. A balance score based on the normalised Shannon entropy is then

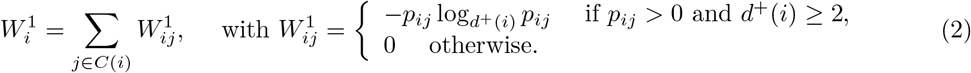

From aforementioned properties of the Shannon entropy, it follows that 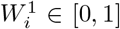 with 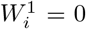 if and only if *d*^+^(*i*) = 1, and 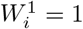 if and only if *i* splits its descendants into at least two equally sized subtrees. Therefore the following specific balance index satisfies our robustness and universality axioms:

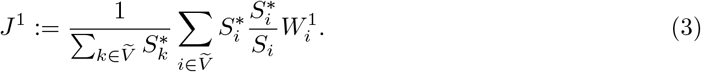

The definition simplifies when we restrict the domain to the set of multifurcating cladograms in which all leaves have equal size *f*_0_ (corresponding to equally important extant types) and internal nodes have zero size (representing extinct ancestors). For all internal nodes *i* in such trees, 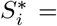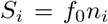, where *n*_*i*_ is the number of leaves of the subtree rooted at *i*. The general definition of Equation 1 then becomes a weighted average of node balance scores:

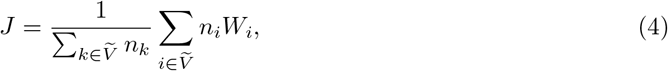

and the specific definition of Equation 3 becomes

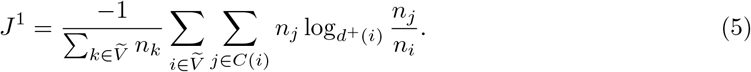

For example, Figure 4 shows the *J*^1^ values of all cladograms on six leaves without linear parts. Unlike *I*_*S*_ and *I*_*C*_, *J*^1^ does not consider the caterpillar tree the least balanced of these cladograms. There are of course many alternative options for *W*. Since the Shannon entropy belongs to a family of generalised entropies ^*q*^*H* (Chao et al., 2014), the above reasoning can be generalised to define a balance score *W*^*q*^, and hence a robust, universal balance index *J*^*q*^, for every *q* > 0 (see Appendix). Other candidates for *W* include one minus the variance of the proportional subtree sizes, or one minus the mean deviation from the median (Mir et al., 2018). We prefer *W*^1^ mostly because, as we shall show, it is the only function for which Equation 4 is a generalisation of the normalised inverse Sackin index.

**Figure 4:**
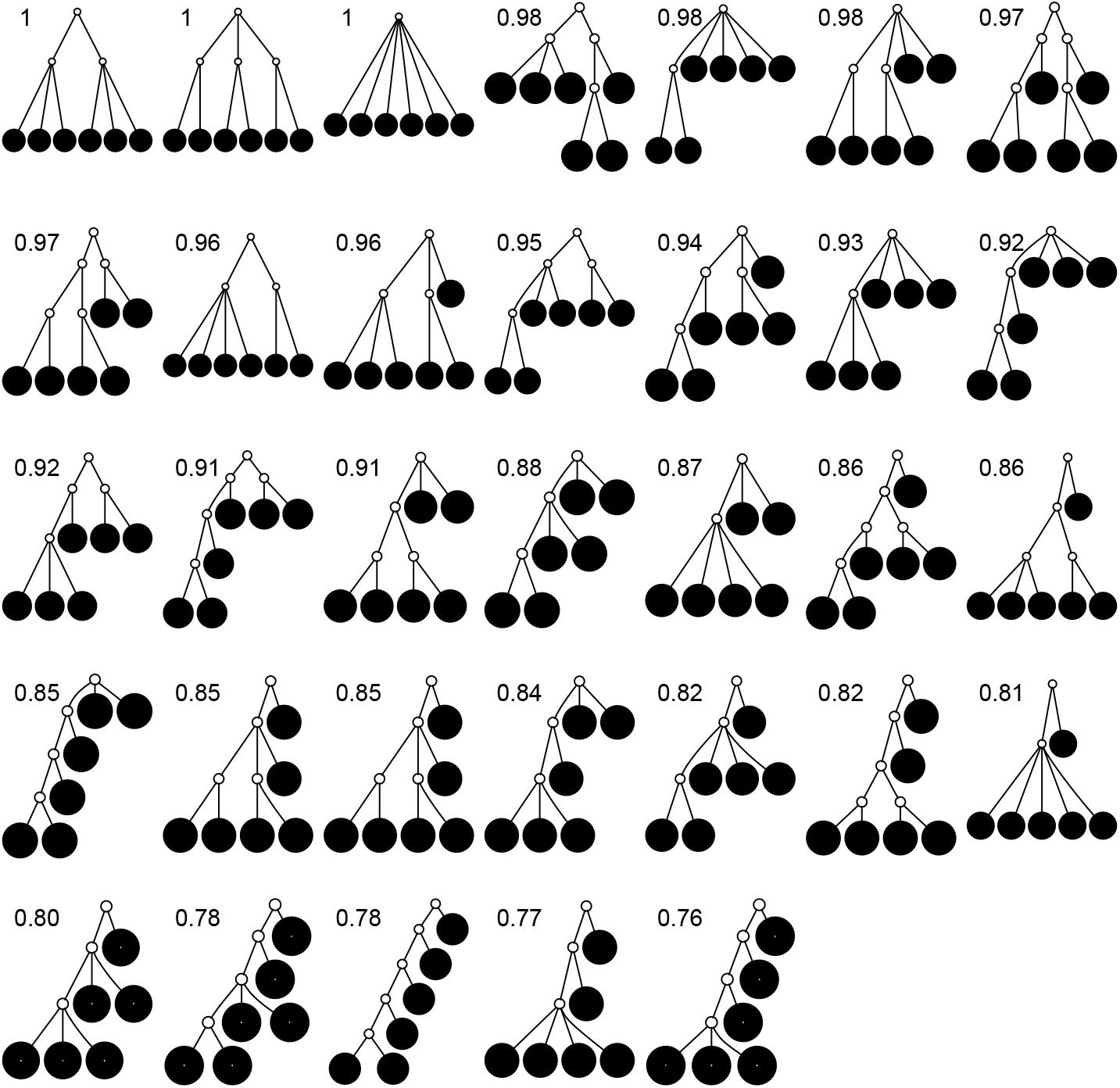
All multifurcating cladograms on six leaves without linear parts and with equally sized leaves, sorted and labelled by *J*^1^ value.

### Relationship with Colless’ index

Like Colless’ index and Colless-like indices as previously defined, our new family of tree balance indices is based on the intuitive idea of assigning a value to each internal node, summing these values, and then normalising the sum. A Colless-like index in the sense of Mir et al. (2018) depends on a function *f* : ℕ → ℝ_≥0_, which assigns node sizes, and a dissimilarity score 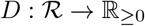, where 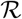 is the set of non-null real vectors. Before normalisation, such an index has the form

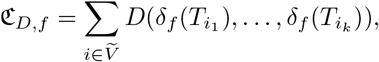

where {*i*_1_, …, *i*_*k*_} are the children of node *i*. The function *δ*_*f*_ assigns a size to each subtree by summing the node sizes: *δ*(*T*) = Σ_*j*∈*V*(*T*)_*f*(*d*^+^(*j*)). Neglecting the initial normalising factor, our general definition (Equation 1) has a similar form and can be considered Colless-like in only a slightly broader sense. Our definition nevertheless differs in three important ways.

First, whereas the unbounded dissimilarity index *D* measures both the relative imbalance of subtrees and their combined size, and is undefined for nodes with out-degree one, we split these two roles into a normalised balance score *W* and an unbounded importance factor *g* and – crucially – we assign a *W* value (specifically zero) to nodes with out-degree one. This difference enables us to extend the balance index definition to trees with any degree distribution. It also makes it easy to normalise our index for any tree, simply by dividing by the sum of the importance factors. Furthermore, our normalisation is universal, rather than being based on comparison with other trees with the same number of leaves. For example, our index judges long caterpillar trees less balanced than short ones (Fig. 5a), whereas Sackin’s index, Colless’ index, and the total cophenetic index consider all caterpillar trees on more than two leaves equally imbalanced.

**Figure 5:**
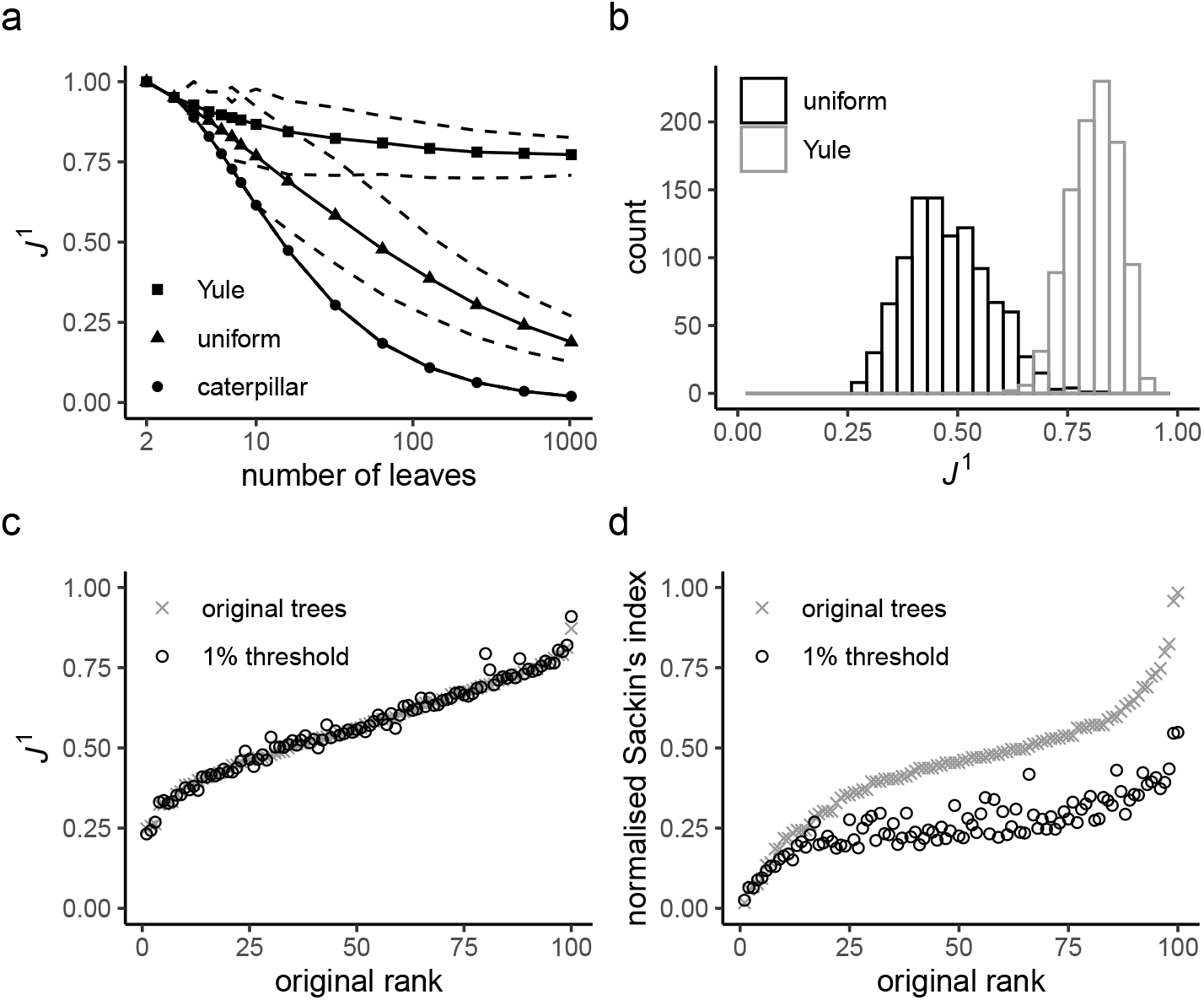
**a**: *J*_1_ values for caterpillar trees and random trees generated from the Yule and uniform models (1,000 trees per data point). All internal nodes have null size and all leaves have equal size. Solid curves are the means and dashed curves are the 5th and 95th percentiles. **b**: *J*^1^ distributions for random trees on 64 leaves generated from the Yule and uniform models (1,000 trees per model). **c**: *J*^1^ values for 100 random trees on 16 leaves, before and after applying a 1% sensitivity threshold. These random trees were generated from the alpha-gamma model with *α* ~ Unif(0, 1) and *γ* ~ Unif(0, *α*). **d**: *I*_*S,norm*_ values for the same set of random trees.

Second, we multiply the balance score by an additional non-root dominance factor *h*. This factor makes the balance index robust when internal nodes can have non-zero size, which blurs the distinction between internal nodes and leaves. Non-root dominance plays no role if all internal nodes have null size, as in cladograms (because then *h* ≡ 1).

Third, instead of assigning a size to each node as a function of its out-degree, we associate a node’s size with the size of the biological population it represents. This ensures that our indices are reliably robust when applied to real data.

### Relationship with Sackin’s index

The sum 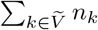 is just another way of expressing Sackin’s index (summing over internal nodes instead of leaves). Therefore *J* in Equation 4 is essentially a weighted Sackin index (with each term in the sum weighted by the balance score *W*) divided by the unweighted Sackin index. In the special, important case of full *m*-ary cladograms, the weighted sum in *J*^1^ (Equation 5) simplifies yet further. Let 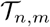 denote the set of all trees on *n* leaves such that all internal nodes have the same out-degree *m* > 1, every internal node has null size, and all leaf sizes are equal. Then we obtain a remarkably simple relationship between *J*^1^ and Sackin’s index:

#### Proposition 0.7.

*Let T be a tree on n leaves with d*^+^(*i*) = *m* > 1 *and f* (*i*) = 0 *for every internal node i. Then*

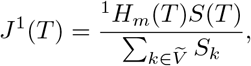

*where* ^1^*H*_*m*_(*T*) *is the Shannon entropy (base m) of the proportional node sizes, and *S*(*T*) is the size of T. If additionally all leaves of T have the same size (so* 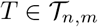 *then*

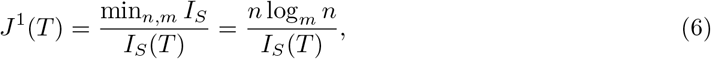

*where* min_*n,m*_ *I*_*S*_ *is the minimum I_S_ value of trees in* 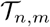.

The above result is somewhat surprising as it unifies our Colless-like index, which can be viewed as a weighted average of internal node balance scores, and Sackin’s index, which is the sum of all leaf depths. A short proof of Proposition 0.7 is presented in the Appendix. The converse result, which is also proved in the Appendix, justifies our choice of *W*^1^ instead of alternative balance score functions:

#### Proposition 0.8.

*Let J be a tree balance index such that*

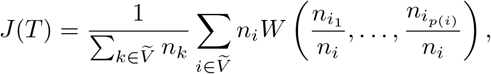

*where i*_1_, …, *i*_*p*(*i*)_ *are the children of node i, and W is a balance score satisfying the conditions stated before Equation 1. Suppose that for all trees* 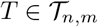, *J*(*T*) = *n* log_*m*_ *n/I*_S_ (*T*). *Then W = W*^1^.

The right-hand side of Equation 6 incidentally provides an alternative way of normalising Sackin’s index on full *m*-ary cladograms, including the bifurcating cladograms on which the index was originally defined. This normalised inverse Sackin index, which we can define as *J*_*S*_ := *n* log_*m*_ *n/I*_S_, provides a more satisfactory way of comparing trees that differ in their node degrees or leaf counts. *J*_*S*_ = 1 if and only if the tree has minimal depth given *m*, which is equivalent to being fully symmetric (because in this case log_*m*_ *n* = *ν*_*l*_ for every leaf *l*). Hence *J*_*S*_ is a *sound* tree balance index in the sense defined by Mir et al. (2018). For *m* > 1, we have *J*_*S*_ > 0 but min *J*_*S*_ 0 as *n*, which makes sense because trees with more leaves can be made less balanced. In particular, when *T* is a caterpillar tree on *n* ≥ 2 leaves,

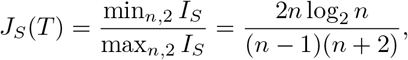

as illustrated in Figure 5a. The definition of *J*_*S*_ can be naturally extended to the case *m* ≤ 1 by setting *J*_*S*_ (*T*) := 0 if *T* is linear or has only one node. From this point of view, *J*^1^ (a Colless-like index) is a generalisation of *J*_*S*_ (the normalised reciprocal of Sackin’s index) to the domain of trees with arbitrary degree distributions and arbitrary node sizes.

### Distributions under the Yule and uniform models

An immediate corollary of Proposition 0.7 is that *J*^1^ can be used to test whether a set of full *m*-ary cladograms is consistent with a particular tree-generating model, with exactly the same sensitivity as Sackin’s index. For example, Figures 5a and 5b show *J*^1^ distributions for random trees generated from the Yule and uniform models, which generate bifurcating cladograms. These two distributions have insignificant overlap when the trees have at least a few dozen leaves.

Kirkpatrick and Slatkin (1993) showed that the expectation of *I*_*S*_ for the Yule model is

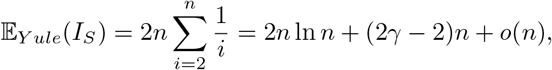

where *γ* is Euler’s constant and *n* is the number of leaves. We find that a good approximation to the *J*^1^ mean for the Yule model is 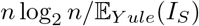, which approaches 1/(2 ln 2) ≈ 0.72 as *n* → ∞.

The expectation of *I*_*S*_ for the uniform model approaches 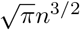 as the number of leaves *n* → ∞ (Blum et al., 2006). Via trial and improvement, we find that a very good approximation to the *J*^1^ mean for the uniform model is *n* log_2_ *n/*(1.7*n*^3/2^ − *n* − 0.808), which approaches zero as *n* → ∞.

### Robustness when applied to random trees

To test the robustness of *J*^1^, we generated random multifurcating trees with node sizes drawn from a continuous uniform distribution, and then compared *J*^1^ values for these trees before and after applying a 1% sensitivity threshold. In the latter case, whenever the combined frequency of a clone and its descendants was below 1%, we merged the corresponding subtree with the clone’s parent, to simulate imperfect detection of rare types. As expected, the *J*^1^ values for the two sets of trees were highly similar, with a median difference of only 0.8% (Figure 5c). In contrast, the median difference in Sackin’s index for the same two sets of trees (after resolving any linear parts in the manner of Figure 2) was 20% (Figure 5d), confirming that *J*^1^ is much more robust than Sackin’s index to the omission of rare types.

### Correlations with preexisting indices

To compare *J*^1^ to Sackin’s index, a Colless-like index, and the total cophenetic index (defined in the Appendix) on a diverse set of trees, we generated 2,000 random multifurcating cladograms on 100 leaves using the alpha-gamma model (Chen et al., 2009) via the R package *CollessLike* (Mir et al., 2018). As shown in Figure 6, our new balance index correlates negatively with the previously defined imbalance indices on this set of random trees, indicating that it captures a similar notion of balance. The strongest correlation is between *J*^1^ and the total cophenetic index (Spearman’s *ρ* = −0.84 for all trees, and *ρ* = −0.97 for trees with mean out-degree greater than 3).

**Figure 6:**
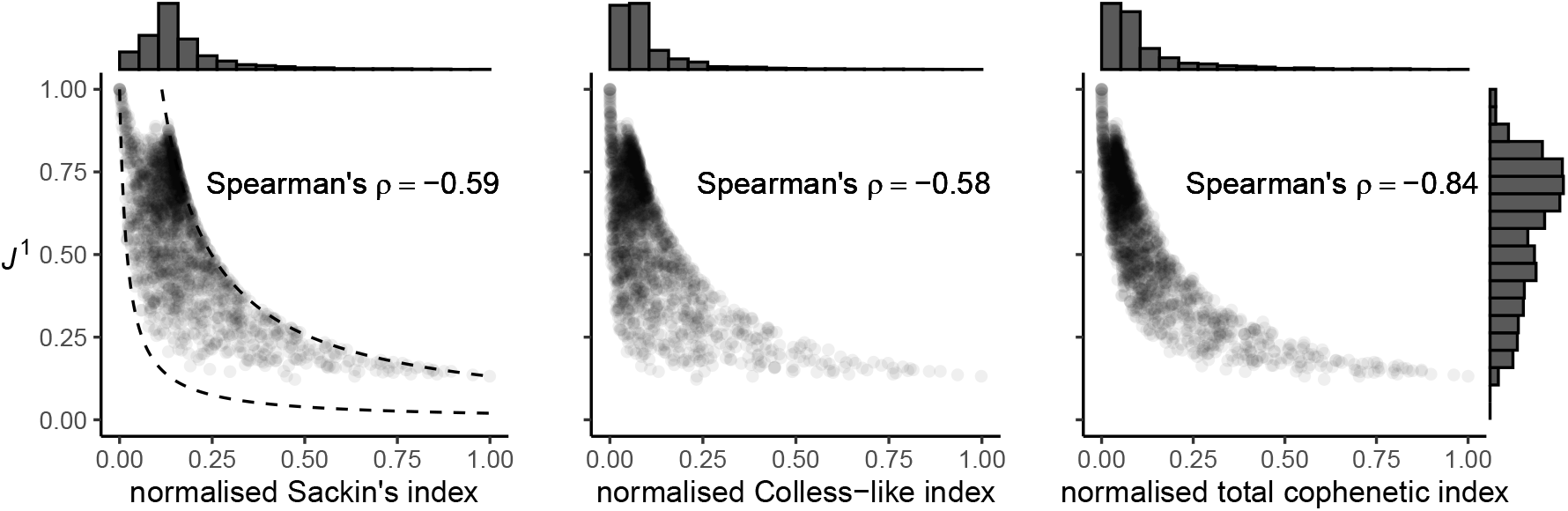
Scatter plots of *J*^1^ versus normalised Sackin’s, Colless-like, and total cophenetic indices for 2,000 random cladograms on 100 leaves. Histograms in the margins show the marginal distributions. Dashed reference curves in the first panel are obtained by substituting *I*_*S,norm*_ into Equation 6 with *n* = 100 and *m* = 2 (upper curve) or *m* = 100 (lower curve). We use the Colless-like index with *f*(*n*) = ln(*n* + *e*) and *D* the mean deviation from the median, as recommended by Mir et al. (2018). Normalisation of each index other than *J*^1^ depends only on the number of leaves and so does not affect correlations. Trees were generated from the alpha-gamma model with *α* ~ Unif(0, 1) and *γ* ~ Unif(0, *α*).

### Discontinuities

Although our indices are robust to the addition of uninformative nodes, the addition of informative nodes – however small – can create a discontinuity. Consider a node that splits its descendants into several subtrees of similar size. The addition of a new, relatively small subtree to this node will create imbalance even as – in fact especially as – the size of this new subtree approaches zero. Our *J*^*q*^ indices are sensitive to this case.

As a more precise example, consider a star tree with *l* > 1 leaves each of size *f*_0_ > 0. Suppose we add to the root another *n* − *l* leaves each of size *x* with 0 ≤ *x* ≤ *f*_0_. The root is assumed to have population 0, so that 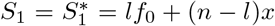 and 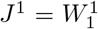. If *x* = 0 then 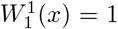 since all *l* leaves have the same size. If *x* > 0 then

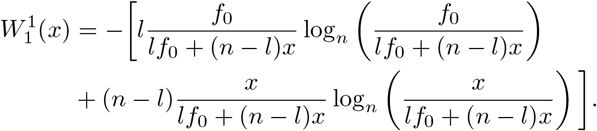

As *x* → 0, so 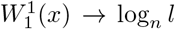 This implies that adding infinitesimally small leaves reduces the balance score from 1 to log_*n*_ *l*, to account for the abrupt loss of balance. The size of the jump is at most 1 − log_3_ 2 ≈ 0.37, and it approaches zero as *l/n* → 1.

### Implementation and algorithmic complexity

Assuming the identity of the root is known, our new indices can be computed from an adjacency matrix in 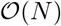 time, where *N* is the number of nodes. Subtree sizes are computed via depth-first search, which takes linear time, and the computation of the balance index takes at most 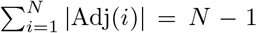 steps, where Adj(*i*) is the adjacency list of node *i*. Efficient R code for calculating *J*^*q*^ is shared in an online repository (Noble and Lemant, 2021).

## Discussion

Here we have defined a new class of tree balance index that unifies, generalises, and in various ways improves upon existing definitions. These indices are applicable to a wider set of trees and enable important new applications.

A challenge in comparing simulated phylogenies and trees inferred from data is that the former are exact, whereas the latter are often incomplete (Scott et al., 2020). In oncology, for example, it has been shown that whether or not a rare tumour clone is detected depends on both methodology and chance (Turajlic et al., 2018). Our balance indices largely solve this problem as they are robust to the omission of rare types, as demonstrated briefly here and more comprehensively in a companion paper (Noble et al., accepted for publication). Besides tumour evolution, our indices are especially well suited to the study of microbial evolution and any other system in which population sizes matter or linear evolution can occur.

In generalising conventional indices we also obviate their shortcomings. Even when restricted to the tree types on which previous indices are defined, our indices enable more meaningful comparison of trees with different degree distributions or different numbers of leaves. This advantage might make our indices preferable to other options more generally.

Because of its unique relationship with Sackin’s index, we especially recommend *J*^1^ – a weighted average of the normalised entropies of the internal nodes – as defined in general by Equation 3 and more simply for cladograms by Equation 5. Given that Sackin’s index has been well studied, it is convenient that *J*^1^ inherits some of the properties of that index when applied to full *m*-ary cladograms, including its relatively high sensitivity in distinguishing between alternative tree-generating models (Kirkpatrick and Slatkin, 1993; Agapow and Purvis, 2002). Within our framework, Sackin’s index is seen not as a general balance index but rather as a normalising factor, which works as a balance index only in the special case of full *m*-ary cladograms (for which the numerator of *J*^1^ is independent of tree topology).

Proposition 0.7 implies that determining the precise moments of *J*^1^ for a model that generates full *m*-ary cladograms is equivalent to determining the moments of the reciprocal of Sackin’s index. Figure 6 suggests that *J*^1^ has interesting relationships with other indices such as the total cophenetic index. These are promising areas for further investigation.

## Acknowledgements

We thank Niko Beerenwinkel, Laura Keller, Francesco Marass, Lisa Lamberti, Jack Kuipers and Katharina Jahn for helpful conversations.

## Author contributions

RN conceived the project. JL and RN developed the balance indices with helpful input from CLS. JL and RN obtained mathematical results with helpful input from VM. JL and RN wrote the paper. All authors have read and approved this manuscript.

## Appendix

## Definition of the total cophenetic index

The cophenetic value *ϕ*(*k, l*) of a pair of leaves (*k, l*) is the depth of their lowest common ancestor. The total cophenetic index (Mir et al., 2013) is then the sum of the cophenetic values over all pairs of leaves:

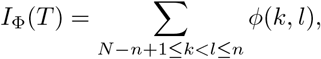

where *N* is the number of nodes and *n* the number of leaves. As in Sackin’s index, the principle is that an unbalanced tree stretches more than a balanced tree. Being explicitly defined for all multifurcating trees, the total cophenetic index permits meaningful comparison of any two multifurcating trees on the same number of leaves.

For trees on *n* > 2 leaves, the minimum of the total cophenetic index is reached on the star tree, with min_*n*_(*I*_Φ_) = 0. The maximum is attained on the caterpillar tree:

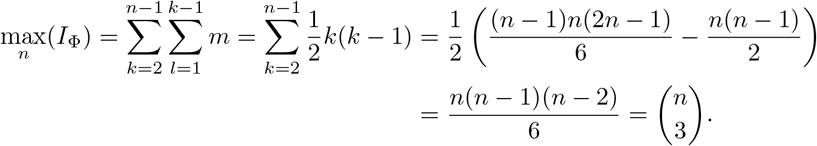

Hence a normalised version of the total cophenetic index is 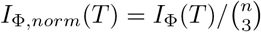. This normalised imbalance index is not minimal for all fully symmetric trees. For example, the cophenetic value of the two leftmost leaves of fully symmetric tree **b** in Figure 1 is two, and so both the unnormalised and normalised cophenetic indices of tree **b** will be nonzero.

## Proof that the index of Equation 1 satisfies our five axioms

*Proof.* Axiom 0.1 (Maximum value): We have *J* ≤ 1 since *h* and *W* lie between zero and one by definition. Also if any internal node *j* of tree *T* doesn’t split its descendants into at least two equally sized subtrees then *W*_*j*_ < 1 by definition and so

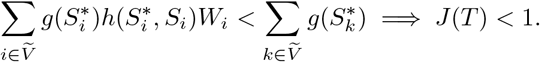

Finally, let *T* be a tree such that every internal node splits its descendants into at least two equally sized subtrees. Then *W*_*i*_ = 1 for all 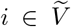 by definition. And if every internal node has null population then 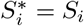, which implies 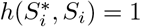 for all 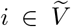 by definition. Hence

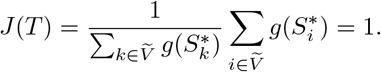

Axiom 0.2 (Minimum value): We have *J* ≥ 0 since *g*, *h* and *W* are always non-negative by definition. Also if *T* is a linear tree then *W*_*i*_ = 0 for all 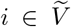 by definition, and hence *J*(*T*) = 0. Conversely, if some internal node *j* has *d*^+^(*i*) > 1 then *W*_*j*_ > 0 by definition and, because *g* and *h* are always positive by definition, we must have *J* (*T*) > 0.

Axiom 0.3 (Leaf limit): Adding a subtree to a leaf *l* changes the tree balance value via the contributions of three sets of nodes: the newly added nodes, the former leaf *l*, and all other internal nodes. First, for each internal node 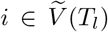 with *i* ≠ *l*, as 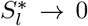 so also 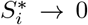 (because 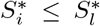), which implies 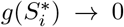 by defination and hence the first contribution approaches zero. Next, as 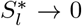, so 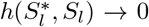 by definition, which implies that the second contribution approaches zero. Lastly, the third contribution approaches zero because *g*, *h* and *W* are continuous by definition.

Axiom 0.4 (Linear limit): Adding a subtree to a node *j*, with previously *d*^+^(*j*) = 1, changes the tree balance value via the contributions of the newly added nodes and of node *j*. The first contribution approaches zero for the same reason as in the leaf limit proof. Now without loss of generality let *j*_1_ denote the original child of *j*, and *j*_2_, …, *j*_*k*_ denote the newly added children of *j*. As 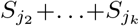 → 0 there are two possibilities. If we also have 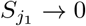 then 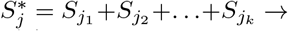 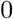, which implies 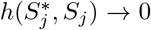 by definition. Otherwise, 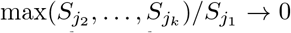, which implies 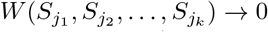 by definition. In either case the second contribution approaches zero.

Axiom 0.5 (Continuity): The continuity of *J* follows immediately from the continuity of *g*, *h* and *W*.

## Other balance indices based on generalised entropies

As defined by Chao et al. (2014), generalised entropies for *q* ≥ 0, *q* ≠ 1 are

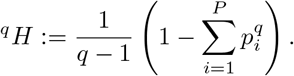

Parameter *q* determines the sensitivity to the type frequencies. ^0^*H* is simply the richness (minus 1) of the population, which corresponds to ignoring the frequencies and just counting the types. For 0 < *q* < 1, rare types are given more weight than implied by their proportion, whereas for *q* > 1 abundant types matter more. ^2^*H* is the Gini-Simpson coefficient. In the limit *q* → 1 we recover the Shannon entropy ^1^*H*_*e*_.

For *q* > 0, ^*q*^*H* attains its maximum value if and only if all types have equal frequency *p*_*i*_ = 1/*m*:

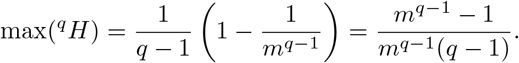

We can therefore define a normalised balance score 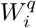 for *q* > 0, *q* ≠ 1 and 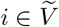 :

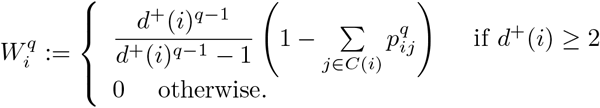

A balance index *J*^*q*^ satisfying our axioms is then

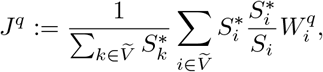

for any *q* > 0. In the limit *q* → 1, *J*^*q*^ → *J*^1^.

## Proof of Proposition 0.7

*Proof.* By definition of *J*^1^, if *T* is a tree on *n* leaves with *d*^+^(*i*) = *m* > 1 and *f*(*i*) = 0 for every internal node *i* then

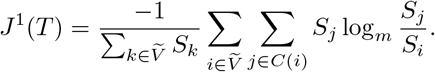

The number of distinct subtrees that contain a given leaf *l* is equal to its number of ancestors, which is the same as *ν*_*l*_, the depth of *l*. So the sum of subtree sizes over the set of all internal nodes is equal to the sum of *ν*_*l*_ multiplied by leaf size over the set of all leaves:

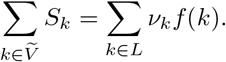

Summing first over the internal nodes and then over their children gives the same result:

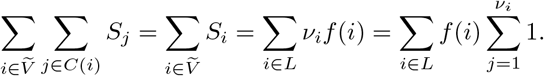

Let *a*(*i, j*) denote the ancestor of node *i* at distance *j*, with *a*(*i*, 0) = *i* and *a*(*i, ν*_*i*_) = *r* (the root) for all *i*. Then by extension,

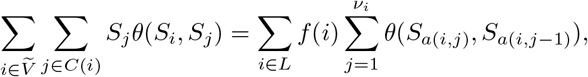

for any function *θ*. In particular, we have

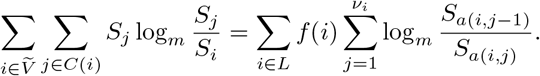

Substituting this result into the expression for *J*^1^ we find

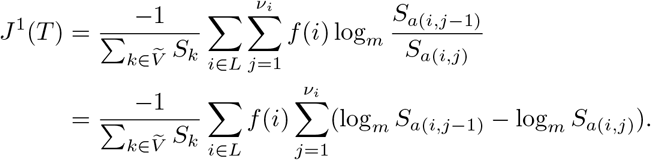

The right-hand sum is a telescoping series that collapses to give

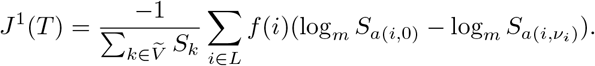

Now since *i* is a leaf, log_*m*_ *S*_*a*(*i*,0)_ = log_*m*_ *S*_*i*_ = log_*m*_ *f*(*i*). Also log_*m*_ *S*_*a*(*i,ν*_*i*_)_ = log_*m*_ *S*_*r*_ = log_*m*_ *S*(*T*). Hence

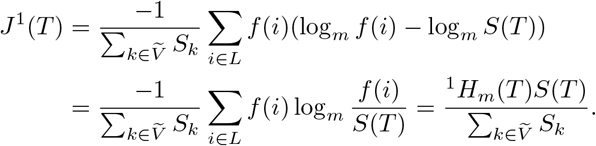

If additionally all leaves *i* of *T* have the same size *f*(*i*) = *f*_0_ then *S*(*T*) = *nf*_0_, ^1^*H*_*m*_(*T*) = log_*m*_ *n*, and 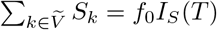 which implies *J*^1^(*T*) = *n*log_*m*_ *n/I*_*S*_(*T*).

## Proof of Proposition 0.8

*Proof.* Since 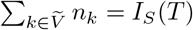 the conditions are equivalent to

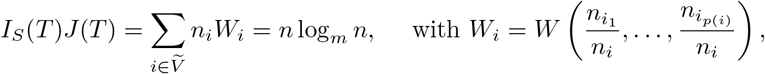

where 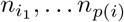 are the children of *i*. Let *T* be a tree in 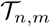 and *i* be an internal node of *T*.

Then 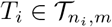 and 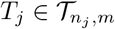 for every child *j* of *i*. Therefore

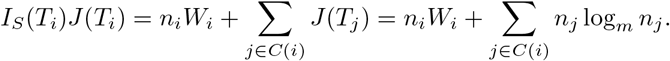

Also, *I*_*S*_(*T*_*i*_)*J*(*T*_*i*_) = *n*_*i*_ log_*m*_ *n*_*i*_, so we have

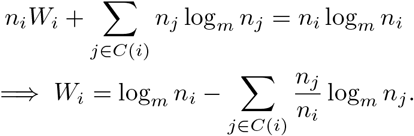

Since Σ_*j*∈*C*(*i*)_ *n*_*j*_ = *n*_*i*_, this implies

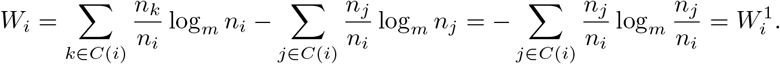

